# A survival-critical role for *Drosophila* giant interneurons during predation

**DOI:** 10.1101/2024.04.30.591368

**Authors:** Cynthia M. Chai, Carmen M. Morrow, Dhyey D. Parikh, Catherine R. von Reyn, Anthony Leonardo, Gwyneth M. Card

## Abstract

Large axon-diameter descending neurons are metabolically costly but transmit information rapidly from sensory neurons in the brain to motor neurons in the nerve cord. They have thus endured as a common feature of escape circuits in many animal species where speed is paramount. Though often considered isolated command neurons triggering fast-reaction-time, all-or-none escape responses, giant neurons are just one of multiple parallel pathways enabling selection between behavioral alternatives. Such degeneracy among escape circuits makes it unclear if and how giant neurons benefit prey fitness. Here we competed *Drosophila melanogaster* flies with genetically-silenced Giant Fibers (GFs) against flies with functional GFs in an arena with wild-caught damselfly predators and find that GF silencing decreases prey survival. Kinematic analysis of damselfly attack trajectories shows that decreased prey survival fitness results from GF-silenced flies failing to escape during predator attack speeds and approach distances that would normally elicit successful escapes. When challenged with a virtual looming predator, fly GFs promote survival by enforcing selection of a short-duration takeoff sequence as opposed to reducing reaction time. Our findings support a role for the GFs in promoting prey survival by influencing action selection as a means to enhance escape performance during realistically complex predation scenarios.

## INTRODUCTION

The large axonal diameter of giant descending interneurons enables rapid information transmission between sensory neurons in the brain and motor neurons via the ventral nerve cord ^1^. Despite the high energetic costs of maintaining such a large cell ^2-4^, giant interneurons are a common feature of escape circuits in many animal species where speed is paramount ^1,5-9^. Although commonly regarded as command neurons that serve as the all-or-none decision-making control element that quickly triggers an escape response ^10,11^, giant neurons are just one of multiple parallel pathways collectively forming a larger escape network ^12-17^. In squid, for example, the interplay between giant and non-giant pathways confers behavioral flexibility during the escape response ^17^. Crayfish can also generate tail flip escape responses via non-giant pathways in response to abrupt tactile threats ^12,14^. Similarly, in flies, genetic silencing of the Giant Fibers (GFs) does not eliminate visually-evoked escape takeoffs, although without GF activity flies only perform longer-duration takeoff sequences that are less likely to launch them off the ground before a theoretical time-of-contact with a virtual stimulus (see also Figure 1A and 1C) ^18^.

**Figure 1.**
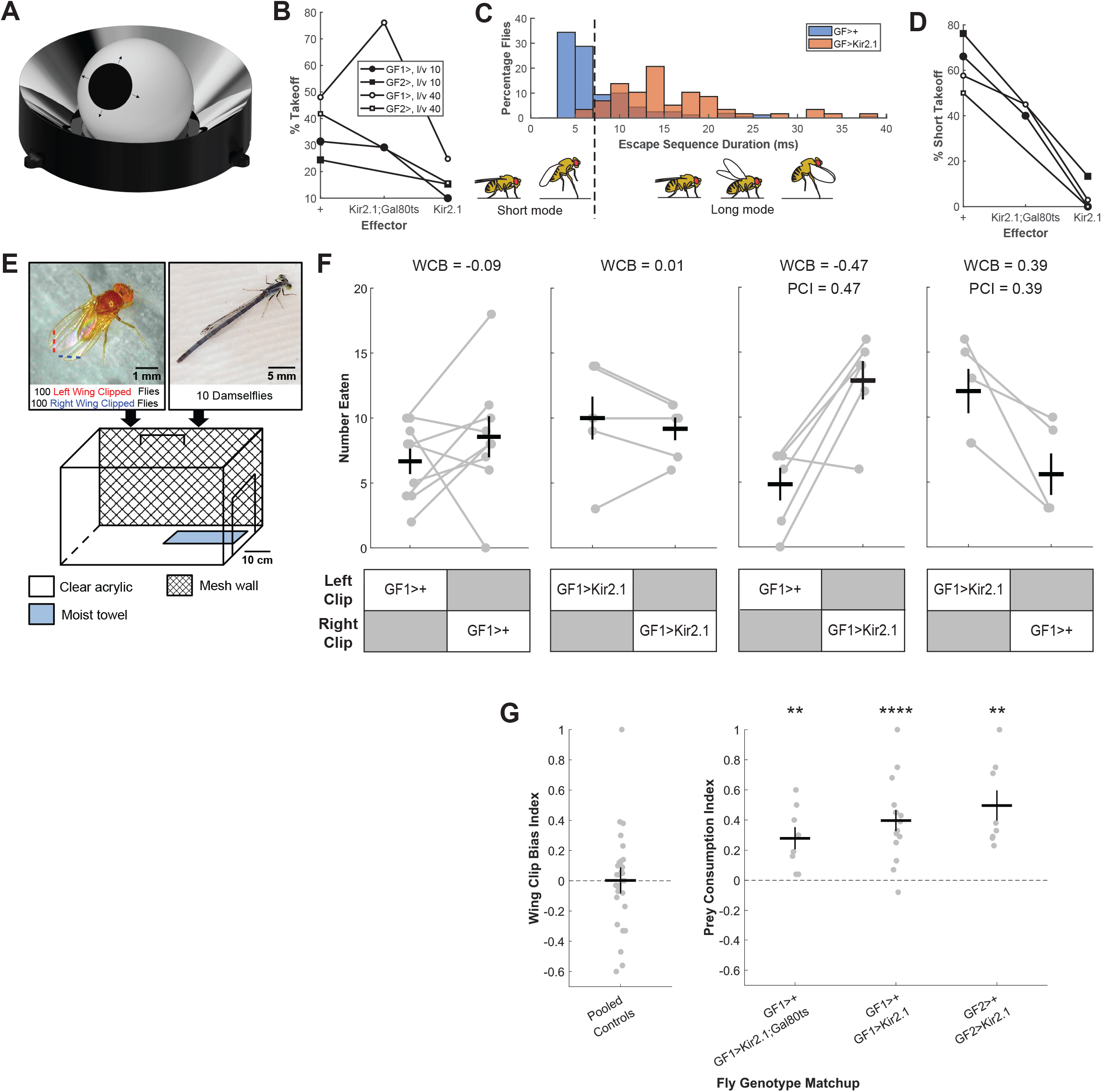
*Drosophila* Giant Fibers promote survival in an ecologically relevant predation regime. (A-D) We tested the effectiveness of genetically silencing the GFs in a virtual predation assay, the FlyPEZ (A), that displays visual looming stimuli to single flies (see Methods). (B) Percentage of flies that took off in the FlyPEZ assay in response to looming stimuli for two different GF-specific split-GAL4 driver line pairings (GF1, GF2) driving expression of the inwardly rectifying potassium channel Kir2.1 to hyperpolarize the GFs and prevent spiking. The Kir2.1; Gal80ts construct provides a weaker silencing effect (see Methods). *l/v*=10: GF1>+ (n=188), GF1>Kir2.1; Gal80ts (n=204), GF1>Kir2.1 (n=139), GF2>+ (n=86), GF2>Kir2.1 (n=98) and *l/v*=40: GF1>+ (n=177), GF1>Kir2.1; Gal80ts (n=222), GF1>Kir2.1 (n=137), GF2>+ (n=134), GF2>Kir2.1 (n=89). n=number of videos analyzed. (C) Histogram of escape sequence durations for flies with functional (GF>+) or silenced (GF>Kir2.1) GFs. The dashed vertical line indicates the intersection of short and long mode distributions. Data are pooled from experiments with both GF1 and GF2 drivers at both fast and slow looming speeds (*l/v*=10, 40; see Methods). GF>+ (n=160) and GF>Kir2.1 (n=58). (D) Percentages of takeoffs that were short mode. Fly genotypes and looming stimulus speeds as in (B). *l/v*=10: GF1>+ (n=59), GF1>Kir2.1; Gal80ts (n=60), GF1>Kir2.1 (n=14), GF2>+ (n=21), GF2>Kir2.1 (n=15) and *l/v*=40: GF1>+ (n=85), GF1>Kir2.1; Gal80ts (n=169), GF1>Kir2.1 (n=34), GF2>+ (n=56), GF2>Kir2.1 (n=14). n=number of videos analyzed. (E) Prey survival competition assay. Each behavior chamber was populated with 10 wild-caught damselflies and 200 flies: 100 with left distal wing tips excised (red dashed line) and 100 with right wing tips excised (blue dashed line). Assays were run for 8 hours. (F) Raw number of flies eaten for a subset of experiments with either the same (WCB) or different genotypes (PCI) in the left and right wing clipped groups. Gray dots represent individual replicates, black crosses represent mean ± SEM. WCB = Wing Clipping Bias index, calculated as the difference in left and right wing-clipped flies eaten divided by the total number of flies (see Methods). PCI = Prey Consumption index, calculated the same as WCB but relative to the GF-silenced vs. control genotypes (see Methods). For both indices, a value of 0 indicates no bias in predation between the competed groups, and a value of ± 1 indicates that only members of one group were consumed. (G) Left - Pooled Wing Clipping Bias index scores for all genotypes (n=30, see also Supplementary Figure 1). Right -Prey Consumption Index scores for between genotype competition trials. Gray dots represent individual trials, black crosses represent mean ± SEM, p-values from one-sample t-test. GF1>+ vs. GF1>Kir2.1; Gal80ts (n=8, **p=0.0068), GF1>+ vs. GF1>Kir2.1 (n=15, ^****^ p=6.0338E-5), GF2>+ vs. GF2>Kir2.1 (n=8, **p=0.0017). n=number of trials.

Depending on the approach speed of an artificial looming visual stimulus, the fruit fly *Drosophila melanogaster* selects between executing a fast escape takeoff motor program or a slower but more stable one ^18^. The switch to a short mode escape depends on GF activation, which can override alternate parallel descending pathways to truncate the escape sequence and propel the fly out of harm’s way once angular size and speed thresholds have been reached ^18^. In the wild, perching flies are at risk from being caught by predators such as the carnivorous odonate, the damselfly ^19,20^. Recordings of damselfly attack trajectories indicate that they attack at speeds near the edge of the fly’s ability to respond. That is, they attack at looming speeds comparable to the fastest virtual stimuli that evoke GF-mediated short mode escapes ^18^. However, the relationship between GF activity and fly escape performance in the context of real predatory danger has not been directly established. Given the degeneracy present in fly escape circuits and the observation that the GF is not necessary for looming-evoked longer-duration takeoffs ^18^, does the GF still confer a survival advantage when a fly is challenged with a real predator?

## RESULTS

### *Drosophila* Giant Fibers promote survival in an ecologically relevant predation regime

We leveraged the rich *Drosophila* transgenic toolkit by driving expression of the inwardly rectifying potassium channel Kir2.1 under the control of GF-specific split-GAL4 drivers (either GF1 or GF2) to achieve constitutive inhibition of only the two bilaterally-paired GF interneurons ^21-23^. To confirm that these transgenic animals had compromised GF function, we used an automated high-throughput behavior system, the FlyPEZ ^24^, to present flies with a virtual looming stimulus that mimics a dark object approaching the fly at either a slow (*l/v*=40) or fast (*l/v*=10) constant velocity (Figure 1A, see Methods). We recorded the time of takeoff for each fly in response to the looming stimulus and calculated the likelihood that flies took off before a hypothetical time-of-contact with the oncoming virtual object. Fewer flies with Kir2.1 silenced GF neurons took off in response to looming compared to split driver-only controls (Figure 1B). For the flies that took off, we measured the duration of the takeoff sequence (time from start of wing raising to time tarsi lost contact with the ground) to determine whether each takeoff was the short (<7 ms) or long (>7 ms) mode. At both slow and fast looming speeds, GF>+ takeoffs were biased towards the faster, short mode escape program (Figure 1C and 1D). However, short mode escapes were almost completely eliminated for GF>Kir2.1 flies (Figure 1C and 1D). Moreover, flies expressing the Kir2.1 effector showed greater short mode escape reduction compared to the weaker Kir2.1; Gal80ts effector, which has a lower UAS copy number (Figure 1D, see Methods). This confirms our previous observations that the GFs drive short mode takeoffs whereas visually mediated long mode escapes are controlled by alternative non-GF neural pathways ^18^.

To determine whether GF-mediated short mode escapes confer a survival advantage under naturalistic predation conditions, we developed a novel prey survival competition assay. In our assay, flies of two different genotypes interacted freely with their natural predators, damselflies, within four custom-built chambers placed in an indoor arena simulating outdoor summer environmental conditions (Figure 1E, Figure S1A). All damselflies used in this study were wild-caught each morning from two sampling sites along a pond within the Janelia Research Campus (Figure S1B). As GF-mediated escape performance does not rely on wing functionality ^25^, fly genotype was marked by excising a small portion of each fly’s wing tip on either the left or right side. Each trial lasted 8 hours, beginning mid-morning and ending in the early evening. At the end of each trial, uneaten flies were anesthetized, sorted based on wing-clipped side, and counted.

As a control, we first competed flies of the same genotype but with different wing sides clipped against each other. A negative wing clipping bias index (WCB) value indicates that more right-wing-clipped flies were eaten compared to left-wing-clipped flies, whereas a positive WCB value indicates that more left-wing-clipped flies were eaten compared to right-wing-clipped flies (Figure 1F, see Methods). The mean WCB values for control experiments with all fly genotypes used in this study were not significantly different from zero indicating that neither right-nor left-wing clipping biased survival chances in this assay (Figure 1G, Figure S1C).

During prey survival competition trials between different fly genotypes, the side of wing clipping (left or right) was alternated between GF-silenced flies and GF driver-only controls to further minimize bias. For each fly genotype matchup, results from competition trials were pooled and a prey consumption index (PCI) calculated (see Methods). A positive PCI indicates that more GF-silenced flies were eaten compared to control flies, with a larger absolute value corresponding to a larger difference between the number of GF-silenced and control flies eaten (Figure 1F). Both GF1>+ (control) versus GF1>Kir2.1 (GF-silenced) and GF2>+ versus GF2>Kir2.1 fly genotype matchups yielded significantly positive mean PCI values indicating that more GF-silenced flies were consumed relative to control animals (Figure 1G). GF1>Kir2.1; Gal80ts flies were also preferentially eaten compared to control flies, though with a slightly smaller mean PCI (Figure 1G). These results establish a direct link between GF activity and fly survival under real predation conditions.

### GF silencing increases likelihood of fly capture

Having established that GF-silenced flies are eaten more frequently than driver-only control flies, we next examined *how* the GFs contributed to survival. To determine whether the greater consumption of GF-silenced flies relative to control animals was directly due to deficiencies in fly escape performance, we video recorded damselfly predation events in a behavior chamber over an 8-hour period (Figure 2A). As fly wing-clipped side, and hence discrimination between different genotypes, was difficult to determine from the video recordings, the chamber was populated by 200 flies (100 left- and 100 right-wing-clipped) of a single genotype during each experiment. Comparing between experiments with different genotypes, we found that the percentage of successful escape outcomes for GF-silenced flies was only half of that for GF driver-only control flies on an individual encounter basis, confirming that the GFs promote fly survival during naturalistic predator attacks, as established in our competition assay results (Figure 2B).

**Figure 2.**
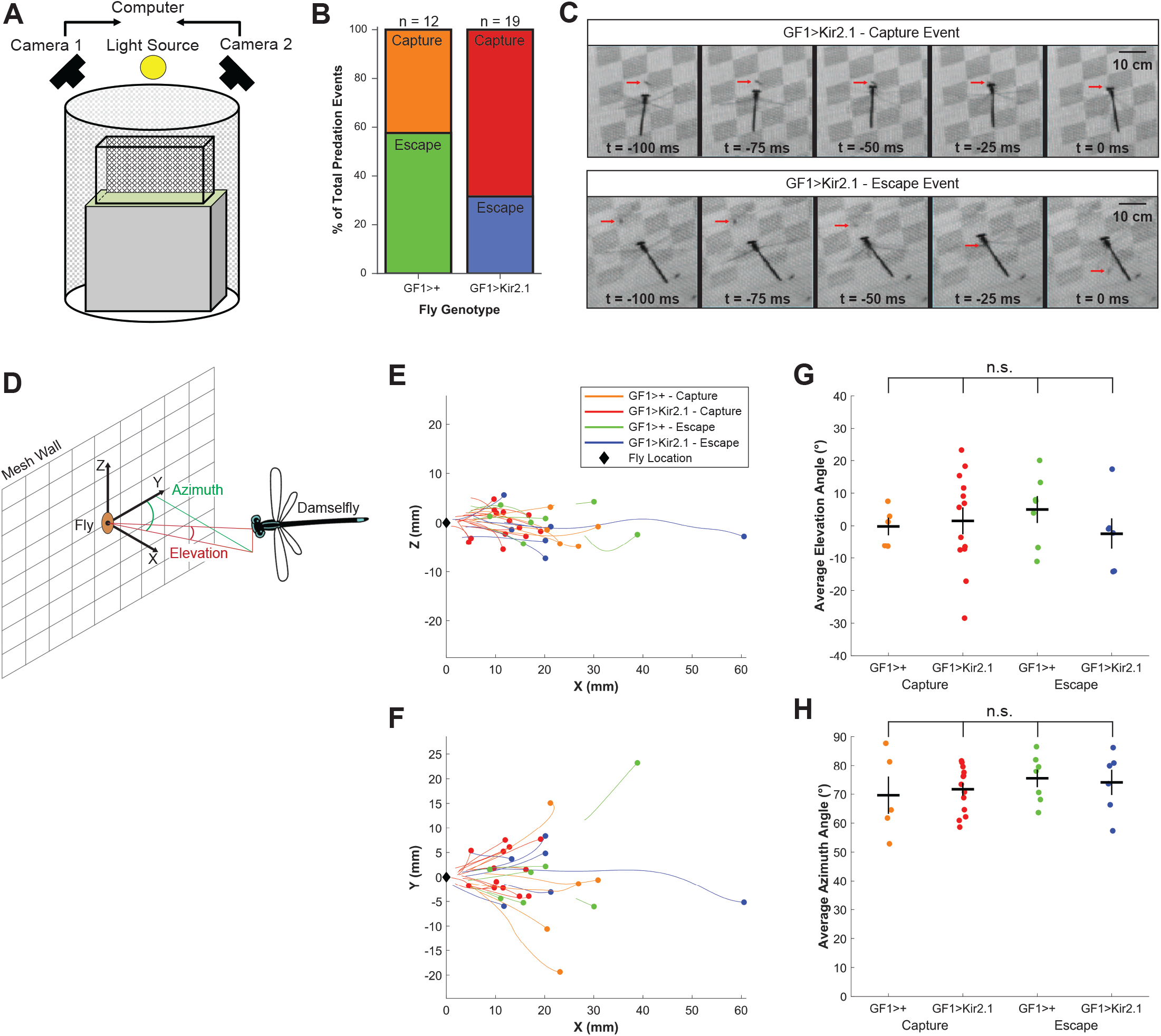
GF silencing increases likelihood of fly capture. (A) Setup of population assay for high-speed videography of predation events. (B) Proportion of recorded predation events that resulted in fly escape or capture outcomes for GF-driver only control and GF-silenced flies. GF1>+ (n=12), GF1>Kir2.1 (n=19). n=number of attack events. (C) Representative video frames showing GF1>Kir2.1 fly capture and escape events. (D) Reference frame coordinates for tracked damselfly attacks. (E-F) Damselfly attack trajectories in the XZ (E) and XY (F) planes. Black diamonds represent damselfly initial positions. Colored dots and lines represent different combinations of fly genotypes and escape outcomes. (G-H) Average damselfly elevation (G) and azimuth (H) angles for each predation attempt. Colored dots represent individual predation events, black crosses represent mean ± SEM. n.s. = not significant, one-way ANOVA followed by Tukey HSD post-hoc test.

We observed that during an aerial attack sequence, damselflies hover at the same height as a fly perching on a wall before darting forward to grab the fly with their legs (Figure 2C). We considered the period after the hover when the damselfly was moving toward the targeted fly as the “attack” and we quantified damselfly attack trajectories by calculating the elevation and azimuth angles of the damselfly relative to the targeted fly during this period (Figure 2D). We found no statistically significant differences in average elevation or azimuth angles over the duration of the attack sequence between prey genotypes (either GF driver-only control or GF-silenced) or capture/escape outcomes (Figure 2E and 2F and 2G and 2H, Figure S2A and S2B). We thus conclude that differences in damselfly attack trajectories were not accountable for the poorer escape performance of GF-silenced flies compared to GF-driver only control animals.

### GF-silenced flies are captured by slower attacks and allow closer damselfly approach

We next analyzed damselfly attack kinematics from the recorded predation events to extract visual features of aerial predator attacks that are selectively detected by the GF ^26-28^. In assays using either GF driver-only control or GF-silenced flies, fly escape outcomes corresponded to damselfly attacks that were approximately constant in approach velocity (Figure 3A). For attacks resulting in control fly capture, however, damselflies accelerated as they approached their prey target with speeds peaking 30 ± 7 ms before time of contact (Figure 3A). A closer look at individual predation events revealed a bimodal distribution in peak speeds and accelerations for GF-silenced capture events (Figure 3B and 3C). While damselfly peak speeds corresponding to control fly capture outcomes were significantly higher than those for escape outcomes, this difference was abolished for GF-silenced flies (Figure 3B). Similarly, peak acceleration values for control capture outcomes were generally higher than escape outcomes while values varied considerably for GF-silenced capture outcomes (Figure 3C). Thus, GF-silenced flies show a consistently reduced ability to escape compared to control flies over ranges of peak speeds and accelerations. These results indicate that GF-silenced flies fail to escape in time when confronted with attack speeds at which they would normally be able to react (Figure 3D, Figure S3A). Interestingly, possibly in response to this prey escape impairment, damselflies in the GF-silenced trial tended to attack at slower speeds compared to the control trial (Figure 3E).

**Figure 3.**
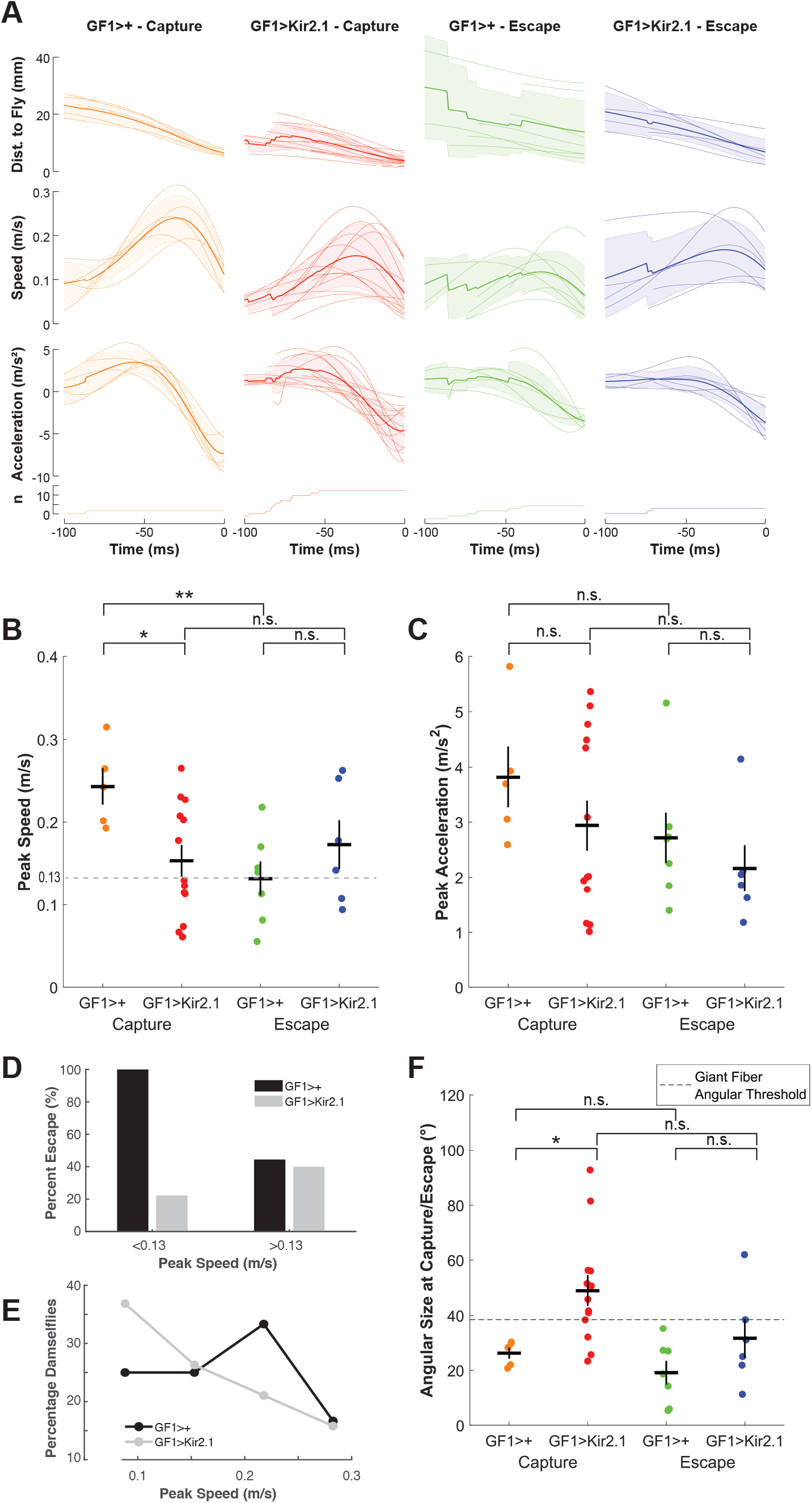
GF-silenced flies are captured by slower attacks and allow closer damselfly approach. (A) Kinematics of damselfly attack trajectories for predation events involving GF-silenced (red, blue) and control (orange, green) flies, clustered by attack outcome (fly capture or escape). Time = 0 represents the time of either contact or escape. Thin lines represent individual attack events and thick lines represent average values. Shaded regions represent the standard deviation. Number of predation events analyzed is indicated in the bottom row. (B) Damselfly peak speeds during attacks. Colored dots represent individual predation events, black crosses represent mean ± SEM. GF1>+ Capture vs GF1>Kir2.1 Capture (*p=0.0176), GF1>+ Capture vs GF1>+ Escape (**p=0.0047), GF1>Kir2.1 Capture vs GF1>Kir2.1 Escape (p=0.5788, n.s.), GF1>+ Escape vs GF1>Kir2.1 Escape (p=0.2677, n.s.). p-values from two-sample t-test. (C) Damselfly peak accelerations during attacks. Colored dots represent individual predation events, black crosses represent mean ± SEM. GF1>+ Capture vs GF1>Kir2.1 Capture (p=0.2989, n.s.), GF1>+ Capture vs GF1>+ Escape (p=0.1525, n.s.), GF1>Kir2.1 Capture vs GF1>Kir2.1 Escape (p=0.3057, n.s.), GF1>+ Escape vs GF1>Kir2.1 Escape (p=0.3987, n.s.). p-values from two-sample t-test. (D) Percentage of GF1>+ and GF1>Kir2.1 flies that escaped damselfly attacks with peak speeds slower or faster than the mean attack speed resulting in escapes for control flies (0.13 m/s, dashed line in Figure 3B). Note the large reduction in GF>Kir2.1 flies that were able to escape the slower attack speeds (<0.13 m/s) that all control flies were able to escape. (E) Distribution of peak attack speeds reached by damselflies during GF1>+ and GF1>Kir2.1 predation events. (F) Angular size of damselfly on fly’s retina at time of contact or escape. Colored dots represent individual predation events, black crosses represent mean ± SEM. GF1>+ Capture vs GF1>Kir2.1 Capture (*p=0.0249), GF1>+ Capture vs GF1>+ Escape (p=0.2197, n.s.), GF1>Kir2.1 Capture vs GF1>Kir2.1 Escape (p=0.0858, n.s.), GF1>+ Escape vs GF1>Kir2.1 Escape (p=0.1488, n.s.). p-values from two-sample t-test. Dashed line indicates the mean angular size at which looming stimuli were previously determined to drive a GF spike and takeoff ^18^.

We observed that damselflies were able to approach more closely to GF-silenced flies, especially for the trials that ended with capture (Figure S3B). Previous studies have shown that a GF spike, and consequently a short mode escape takeoff, is triggered once a looming stimulus reaches a threshold angular size of approximately 40° on the fly’s retina ^18^. We examined the relationship between the maximum angular size a damselfly subtended at its closest distance to the fly and fly escape outcomes for both genotypes. To do this we used the damselfly head width (Figure S3C), the widest part of the damselfly body from the fly’s perspective during a direct attack, to convert the distance of the damselfly at the point of fly capture or escape to an estimated angular size that the damselfly would subtend on the fly’s retina (Figure 3F). Damselflies for GF-silenced fly capture outcomes were the only ones that consistently and considerably surpassed the GF activation threshold (Figure 3F). This implies that a role of the GF is to ensure the fly escapes before a predator comes close enough to capture it with high probability. Furthermore, the observation that the GF’s 40° threshold is consistently surpassed in GF-silenced flies indicates that this is a unique role for this single pair of giant axon descending neurons – there is no other fail safe or redundant circuit for this function.

### GF increases escape likelihood by promoting the shorter-duration escape takeoff sequence

The GFs triggering escape at a certain size threshold as a way to prevent predators from getting too close is at odds with the idea that giant axons facilitate fast transmission enabling the animal to reduce their reaction time. To directly assess whether fly reaction times were altered by GF silencing, we reanalyzed the behavioral responses of GF-driver only control and GF-silenced flies that were challenged with virtual looming stimuli in the FlyPEZ (Figure 4A). We manually annotated the frame in which the escape takeoff sequence started (when the fly’s wing first begins to elevate) and when the sequence concluded (when the first tarsus of the two middle legs leaves the ground) (Figure 4B). We consider the reaction time to be the former (earlier) of these two time points, and the escape time to be latter (later) of these two time points. We found that the range of reaction times was quite large and that there was no significant difference in median reaction time between GF-silenced and GF driver-only control flies for either the slow (*l/v*=40) or fast (*l/v*=10) stimulus (Figure 4C). When we examined the timing of the end of the escape sequence when the fly lifts off, however, we found that escape time was significantly shorter for control compared to GF-silenced flies in the case of the faster looming stimuli (*l/v*=10) (Figure 4D). This indicates that the functional consequence of our earlier observation that GF driven escape sequences are shorter than non-GF ones (Figure 1C), is that even without reacting sooner, flies can still escape quicker, and that this is likely relevant to surviving naturalistic damselfly attacks. Taken together, our results argue that the role of the *Drosophila* GFs is not to reduce reaction time during a predator attack but rather to mediate action selection, ensuring the fly elects to use its short-mode takeoff sequence to get off the ground in time if and when a looming threat gets close enough that capture is imminent.

**Figure 4.**
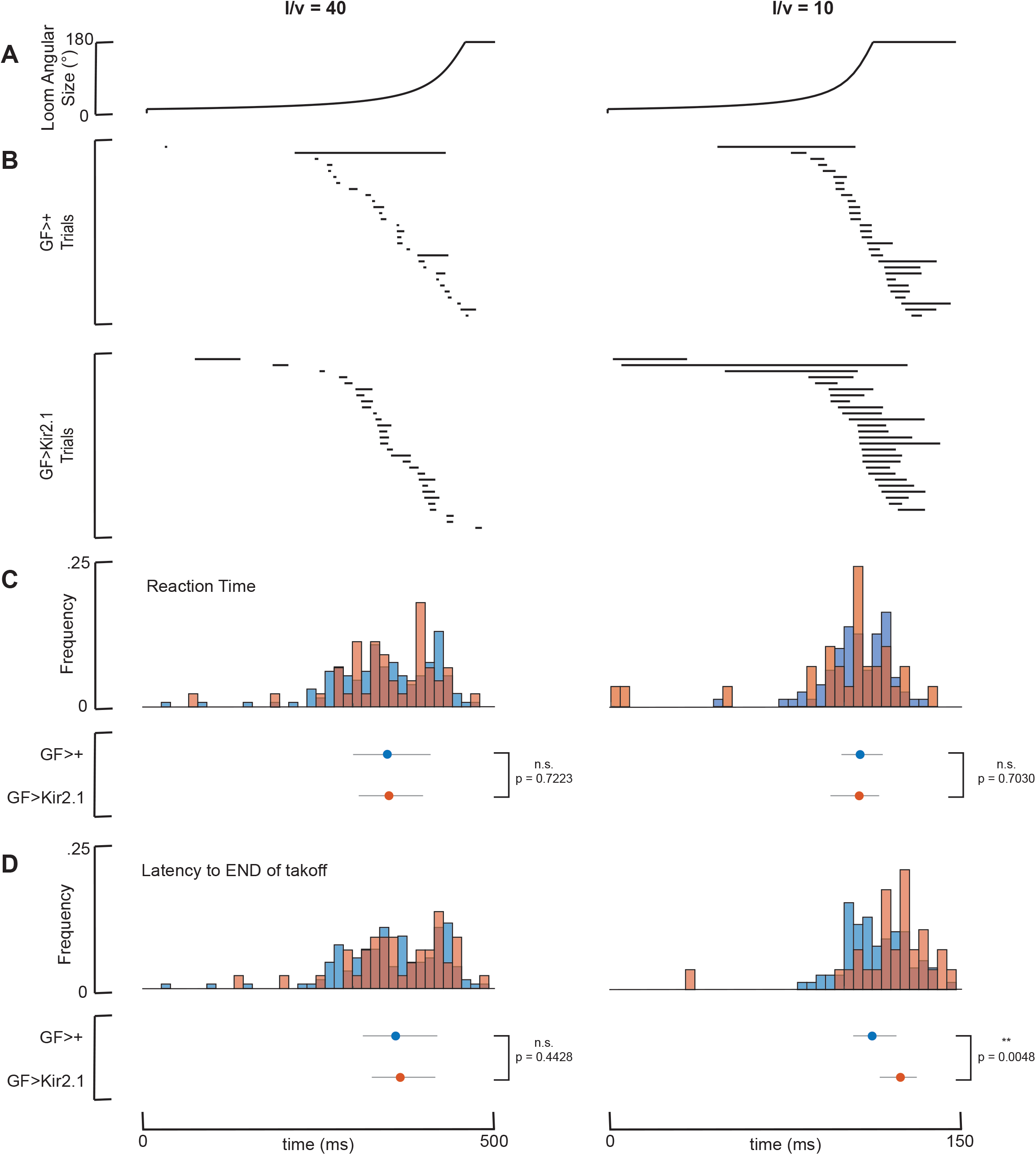
GF increases escape likelihood by promoting the shorter-duration escape takeoff sequence. (A-D) GF>+ and GF>Kir2.1 fly escape sequences for *l/v*=40 (left column) and *l/v*=10 (right column) loom expansion speeds in FlyPEZ assays. Note shorter time axis range for *l/v*=10 column. (A) Time course of looming stimulus angular size expansion. (B) Takeoff sequence ethograms. Dark lines mark the duration of the takeoff sequence, from wing lift initiation to loss of ground contact for representative individual flies, n=29 flies per genotype per loom expansion speed. (C) Histogram (above) and median ± interquartile range (below) of reaction time (latency to start of takeoff sequence). Data are pooled from experiments with both GF1 and GF2 drivers. GF>+, *l/v*=40 (n=141), GF>Kir2.1, *l/v*=40 (n=48), GF>+, *l/v*=10 (n=80), GF>Kir2.1, *l/v*=10 (n=29). n.s. = no significance, two-sample t-test. (D) Histogram (above) and median ± interquartile range (below) of latency to the *end* of the takeoff sequence. Data are pooled from experiments with both GF1 and GF2 drivers. GF>+, *l/v*=40 (n=141), GF>Kir2.1, *l/v*=40 (n=48), GF>+, *l/v*=10 (n=80), GF>Kir2.1, *l/v*=10 (n=29). n.s. = no significance, **p < 0.01, two-sample t-test.

## DISCUSSION

Over the course of evolution, predation risk in the wild exerts large selection pressures on the form and function of prey escape circuits. A hallmark of escape circuit architecture is degeneracy – in crayfish, teleost fish, and flies, both giant and non-giant neuron-mediated pathways connect sensory detection of a predator to motor neurons coordinating an escape ^14,18,29^. The ability to recruit multiple parallel pathways in different combinatorial patterns lends flexibility to these rapid responses, enabling fine-tuning of the escape response to each predation scenario. However, the metabolic cost of maintaining a neuron increases proportional to the neuron’s volume ^2-4^. Including a giant neuron in a degenerate network should thus be selected against unless its presence somehow increases the animal’s fitness beyond the contributions of non-giant circuits.

Despite the evolutionary significance of predator evasion responses, few attempts have been made to determine the survival value and role of giant interneurons under ecologically relevant conditions. Two decades ago, giant-independent crayfish tail flips were shown to be less effective than giant-mediated ones at preventing capture during dragonfly nymph attacks ^30^. A more recent study in zebrafish also confirmed the life preserving role of the giant Mauthner cell when subjected to damselfly nymph predation ^31^. Both these studies demonstrated that the role of the giant neurons in these systems was to reduce the time between predator detection and the first escape action, improving the prey’s likelihood of escaping before capture. This behavioral role for giant neurons matches their biophysical adaptation of enlarged axon diameter, which increases action potential speed by decreasing the axial resistance of the main neuron cable ^32^. Thus, it has been presumed that the role of giant neurons in escape circuits is to reduce prey reaction time.

Our results challenge this narrative as the sole explanation for giant neurons in escape circuits, as we find that *Drosophila* GFs play a different role. We find that GF silencing does not alter reaction time, measured as the latency to when the takeoff sequence *starts*. Instead, the GF’s enforce selection of a specific short-duration escape takeoff sequence that reduces the latency to when the takeoff sequence *ends*. In this context, the transmission speed conferred by the large diameter of the GF axon performs an action selection role that short circuits parallel pathways that would drive longer escape takeoff sequences, enforcing use of the shorter, life-saving action.

Previous circuit-level analysis show that the GF integrates input from two different types of visual projection neurons, LPLC2 and LC4 ^26-28^. These neurons provide information about the angular size and expansion velocity of looming stimuli, respectively, to trigger a short mode escape when a rapidly expanding virtual looming stimulus reaches about 40° on the fly’s eye ^27,28^. Our findings also support this circuit model at a behavioral ecology level. We show that the GFs are necessary for flies to consistently escape before an attacking damselfly’s head reaches a size of 40° on its retina. This critical size of 40° corresponds to when the damselfly is about 3 mm away from the fly, which may be the edge of the range at which the damselfly can comfortably extend its four front legs to grab the fly. We note that in our observed predator-prey interactions with GF-silenced flies, the fly was captured in all but one of the events in which the damselfly approached GF-silenced flies more closely than this GF angular threshold.

By combining approaches in behavioral neuroscience and ecology, we provide experimental evidence that the GFs promote prey fitness by enhancing fly escape performance when challenged with damselfly attacks in a naturalistic setting. Our analysis of damselfly attack kinematics further indicates that the GF generates timely escape responses by integrating multiple predatory visual features including attack speed and angular size of the predator on the fly’s retina. We conclude that despite the presence of parallel escape pathways, GF-mediated fast escapes are still strongly accountable for successful predator evasion and survival under ecologically relevant conditions.

## AUTHOR CONTRIBUTIONS

Conceptualization: G.M.C., C.M.C., C.R.v.R., and A.L.; Methodology: G.M.C., C.M.C., C.M.M., and A. L.; Investigation: C.M.C.; Data curation: C.M.C.; Formal analysis: C.M.M., C.M.C., and D.P.P.; Software: C.M.M.; Visualization: C.M.M. and C.M.C.; Writing – Original draft: C.M.C.; Writing – Review and editing: C.M.C., C.M.M., and G.M.C.; Funding acquisition: G.M.C.; Resources: G.M.C. and A.L.; Project administration: G.M.C.; Supervision: G.M.C.

## ACKNOWLEDGMENTS

We thank Grace Zheng (Janelia FlyCore) for assistance with fly husbandry, Huai-Ti Lin for advice on high-speed videography, the Janelia Instrument Design and Fabrication Team for behavior chamber construction, Card lab members for helpful comments on the manuscript, and Alison Howard for administrative assistance.

## MATERIALS AND METHODS

### EXPERIMENTAL MODEL AND SUBJECT DETAILS

We used 2 to 5-day-old female *Drosophila melanogaster* reared on standard cornmeal fly food at 22-25**°**C and 50% humidity with a 16-h light/8-h dark cycle. The following fly stocks were used in this study: *DL* (“+”, wild-type strain from M. H. Dickinson), *split-GAL4(R17A04_p65ADZp (attP40); R68A06_ZpGdbd (attP2))* (GF1), *split-GAL4(R14A01_p65ADZp (attP40); R79H02_ZpGdbd (attP2))* (GF2), *w+;; pJFRC49-10XUAS-IVS-eGFPKir2*.*1 (attP2)* (Kir2.1), *w+; tubP-Gal80ts; UAS-Kir2*.*1* (Kir2.1;Gal80ts). The *tubP-Gal80ts* transgene is not functional when crossed with split-GAL4 driver lines such as GF1 and GF2 ^33^. The *pJFRC49-10XUAS-IVS-eGFPKir2*.*1* construct has a higher UAS copy number compared to the *tubP-Gal80ts; UAS-Kir2*.*1* construct and should produce a stronger neuronal silencing effect.

The day before prey survival competition and predation high-speed videography experiments, flies were cold anesthetized for a maximum of 20 minutes while either their left or right wing tips were excised using dissection scissors. Male and female damselflies belonging to the genus *Ischnura* were wild-caught from a pond within the Janelia Research Campus in Ashburn, Virginia, USA using a butterfly net and kept in 100 mm plastic Petri dishes on ice 1-2 h before experiments. All experiments were conducted between the months of July and September.

### METHOD DETAILS

#### FlyPEZ Assay

We used a high-throughput behavioral assay, FlyPEZ, to record the responses of unrestrained flies presented with computer-generated looming stimuli ^24^. In accordance with a previous study ^34^, we characterized the looming time course by the size-to-speed ratio, *l/v*, of the virtual object, where *l* = virtual object radius and *v* = virtual object approach velocity. Briefly, individual flies were released through an automated gate onto a prism where they were presented with a dark disk stimulus projected onto a white dome. The dark disk expanded to a fixed size at different looming rates characterized by their *l/v* ratio (fast looming, *l/v*=10 ms; slow looming, *l/v*=40 ms). If flies failed to takeoff upon stimulus presentation, the stage was cleared before release of the subsequent fly. All experiments were conducted during the four-hour time window before incubator lights were switched off, which coincides with the flies’ evening activity peak. A single stimulus was presented per fly, and videos were recorded at 6000 frames per second under 740 nm infrared illumination. To quantify the duration of the escape takeoff motor sequence, videos were manually annotated for the start (first frame of wing raising) and end (first frame when the two middle “jumping” legs left the ground) of the escape sequence. A total of 1474 flies were screened. For Figure 1C, to minimize bias towards *l/v*=10 or *l/v*=40 data after pooling, the n in the larger group was decreased by randomly selecting points to remove in order to match the n in the smaller group.

#### Prey Survival Competition Assays

##### Experimental Design

For the prey survival competition assay, we released 100 GF>+ flies (GF split-GAL4 driver lines crossed to DL), 100 GF>Kir2.1 or GF>Kir2.1; Gal80ts (GF split-GAL4 driver lines crossed to either Kir2.1 or Kir2.1; Gal80ts) flies, and 10 damselflies into a 15 x 15 x 20 cm clear plastic behavior chamber. For wing clipping bias assays, we released 100 left wing clipped and 100 right wing clipped flies of the same genotype into the chamber with 10 damselflies. Each behavior chamber had a mesh wall on the long face to facilitate fly perching and the equilibration of internal and external conditions. A wet paper towel was taped down to the floor of each chamber at the start of each assay period to minimize animal dehydration.

##### Experimental Conditions

To simulate naturalistic conditions, we placed the boxes inside an environmental room that was illuminated to outdoor conditions and wallpapered with naturalistic scenery. Temperatures in the room cycled between 31**°**C during the day and 19**°**C overnight. Four behavior chambers were assayed simultaneously during each 8-hour assay period, which began at around 10 am and ended at around 6 pm.

##### Fly Collection and Counting

At the end of each experimental period, flies were anesthetized using FlyNap (a mixture of 50% triethylamine, 25% fragrance, 22.63% ethanol, 1.25% 2-propanol, and 1.13% methanol purchased from the Carolina Biological Supply Company). Anesthetized flies from each chamber were then collected in a flask connected to a vacuum line and left in the refrigerator for a few hours before counting. The wing clipping bias index (WCB) and prey consumption index (PCI) were calculated using the following formulas:

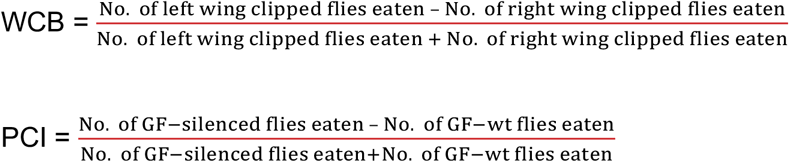

#### High-speed Videography of Damselfly Predation Events

Similar to the wing clipping bias assays, a behavior chamber containing 100 left wing clipped and 100 right wing clipped flies of the same genotype were released into a chamber with 10 damselflies. The chamber was placed on a podium covered with artificial green turf within a cylindrical enclosure lined with checkered wallpaper to provide optic flow. Two high-speed cameras (Photron SA-1) were positioned orthogonal to each other around the enclosure with both cameras focused on the chamber’s mesh wall. Assays were performed outside the indoor environmental arena at room temperature to facilitate video recording. Throughout an 8-hour assay period, an experimenter observed the cameras’ live feeds on a PC monitor and manually triggered the cameras to record predation events. Videos were recorded at 1,000 frames per second using Photron software. GF-wild type and GF-silenced flies were assayed separately on consecutive days. After video acquisition, videos were screened for true predation attempts using the following criteria: 1) the damselfly hovers in place behind the perching fly during attack sequence initiation, 2) moves forward towards target, and 3) forms a basket with its legs to grab the fly. If any one of these three criteria were not met, the video was discarded. An attack event was scored as a capture if the damselfly grabbed the fly off the mesh wall. An attack event was scored as an escape if the fly jumped away from the wall before the damselfly could reach it. Flies that were initially caught but eventually freed themselves from the damselflies’ grasp were still scored as being captured.

#### Predation Video Digitization and 3D Kinematic Analysis

Damselfly head, damselfly tail, and fly body centroid points were marked in each video frame using DLTdv software ^35^. Damselfly attack trajectories were analyzed using custom scripts in MATLAB. Each component axis was passed through a Butterworth filter followed by a Savitzky-Golay filter and attack kinematics parameters were calculated. Due to left-right symmetry, the absolute values of the azimuthal plane coordinates (XY) were used to calculate average trajectory azimuth angle, constraining the angle between 0° and 180°. The angular size of the damselfly on the fly’s retina at time of escape or capture (θ) was calculated using the formula θ = 2tan^-1^(*r*/*d*) where 2*r* is the damselfly head width and *d* is the Euclidean distance between the damselfly and the fly at time of escape or capture ^18^.

### QUANTIFICATION AND STATISTICAL ANALYSIS

For prey survival competition assay (Figure 1, Figure S1), student’s one sample t-test was used to determine statistically significant differences from zero for prey consumption index and wing clipping bias index scores. For damselfly attack elevation and azimuth analysis (Figure 2), one-way ANOVA followed by Tukey HSD post-hoc test was used to determine statistically significant differences between all groups. For damselfly attack kinematics analysis (Figure 3, Figure S3), student’s two-sample t-test was used to determine statistically significant differences between groups of interest. For FlyPEZ assays (Figure 4), student’s two sample t-test was used to determine statistically significant differences between fly genotypes. Any additional information including custom MATLAB codes required to reanalyze the data reported in this paper is available from the lead contact upon request.

## SUPPLEMENTAL FIGURES

**Figure S1.**
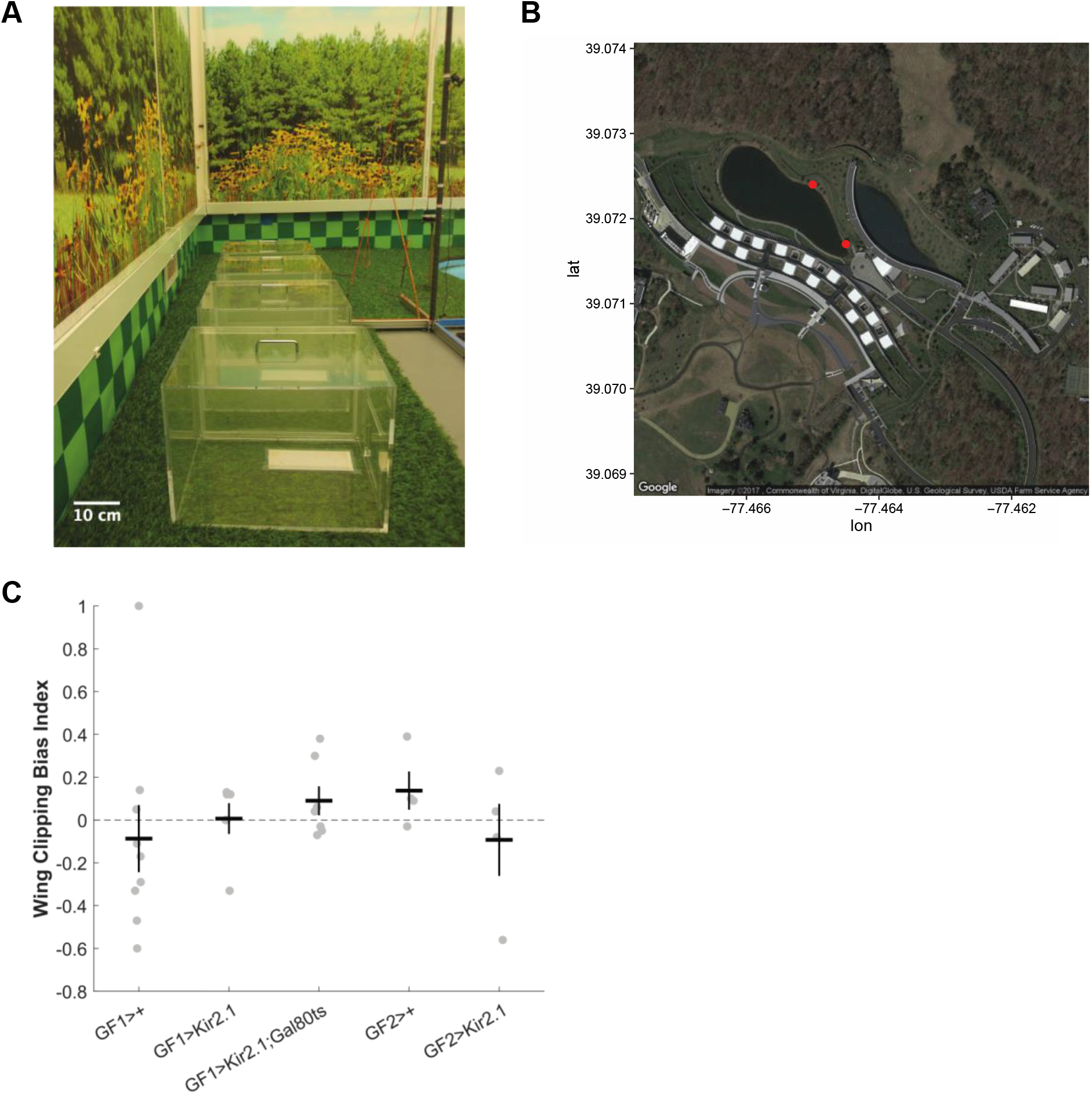
Prey survival competition assay. Related to Figure 1. (A) Four behavior chambers assayed simultaneously inside an indoor arena that simulates outdoor summer environmental conditions. (B) Damselfly collection sites (red dots) within the Janelia Research Campus, Ashburn, VA. Map generated using ggmap package in R. (C) Wing clipping bias index scores for within genotype controls. Gray dots represent individual trials, black crosses represent mean ± SEM, p-values from one-sample t-test. GF1>+ (n=9, p=0.5955), GF1>Kir2.1 (n=6, p=0.9296), GF1>Kir2.1;Gal80ts (n=7, p=0.2306), GF2>+ (n=4, p=0.2209). GF2>Kir2.1 (n=4, p=0.6210). n=number of trials.

**Figure S2.**
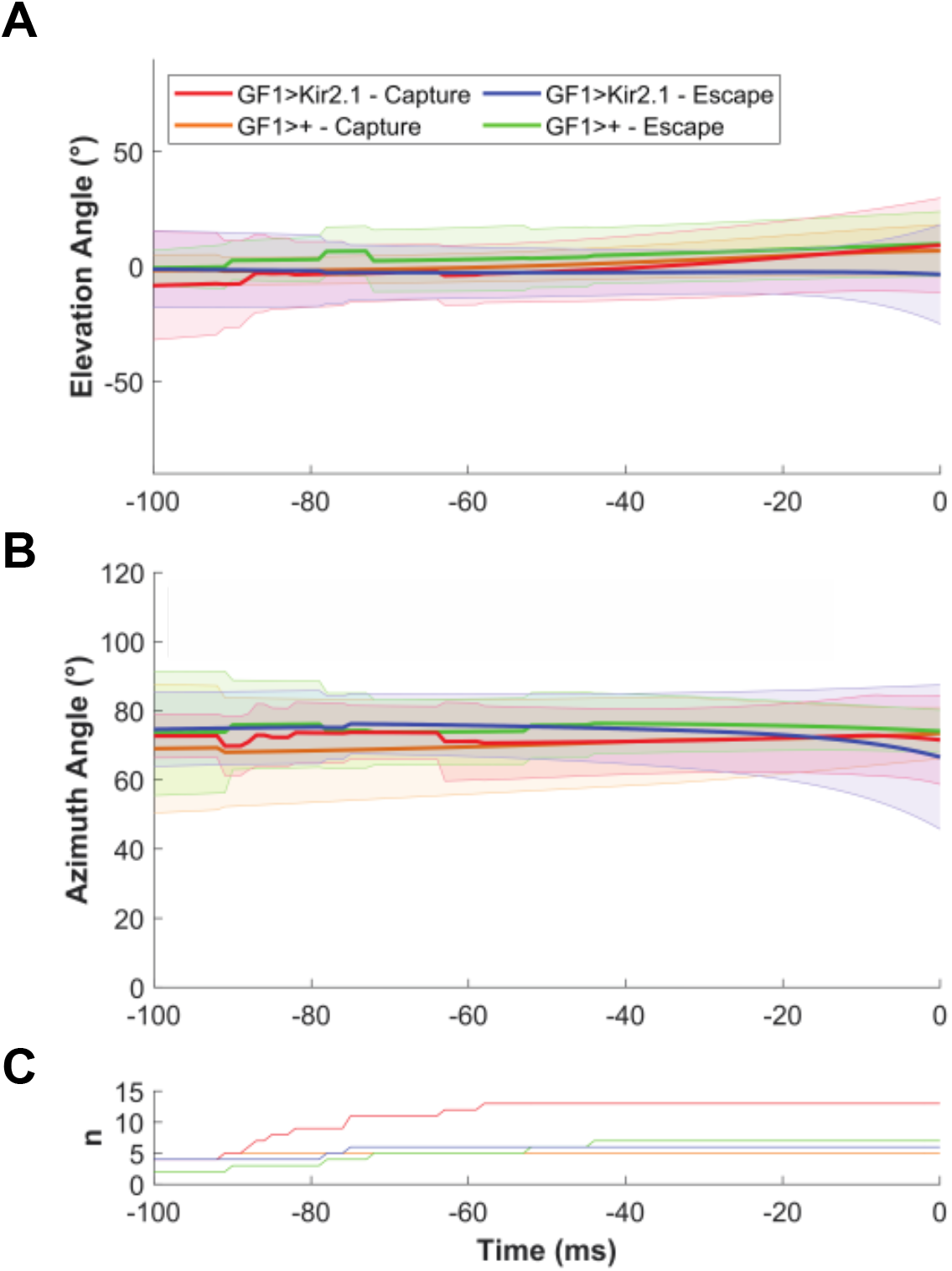
Time course of damselfly elevation and azimuth angles during attack events. Related to Figure 2. (A-C) Average elevation (A) and azimuth (B) angles and number of traces averaged (C) as functions of time. Time = 0 represents the time of either contact or escape. Shaded regions represent the standard deviations.

**Figure S3.**
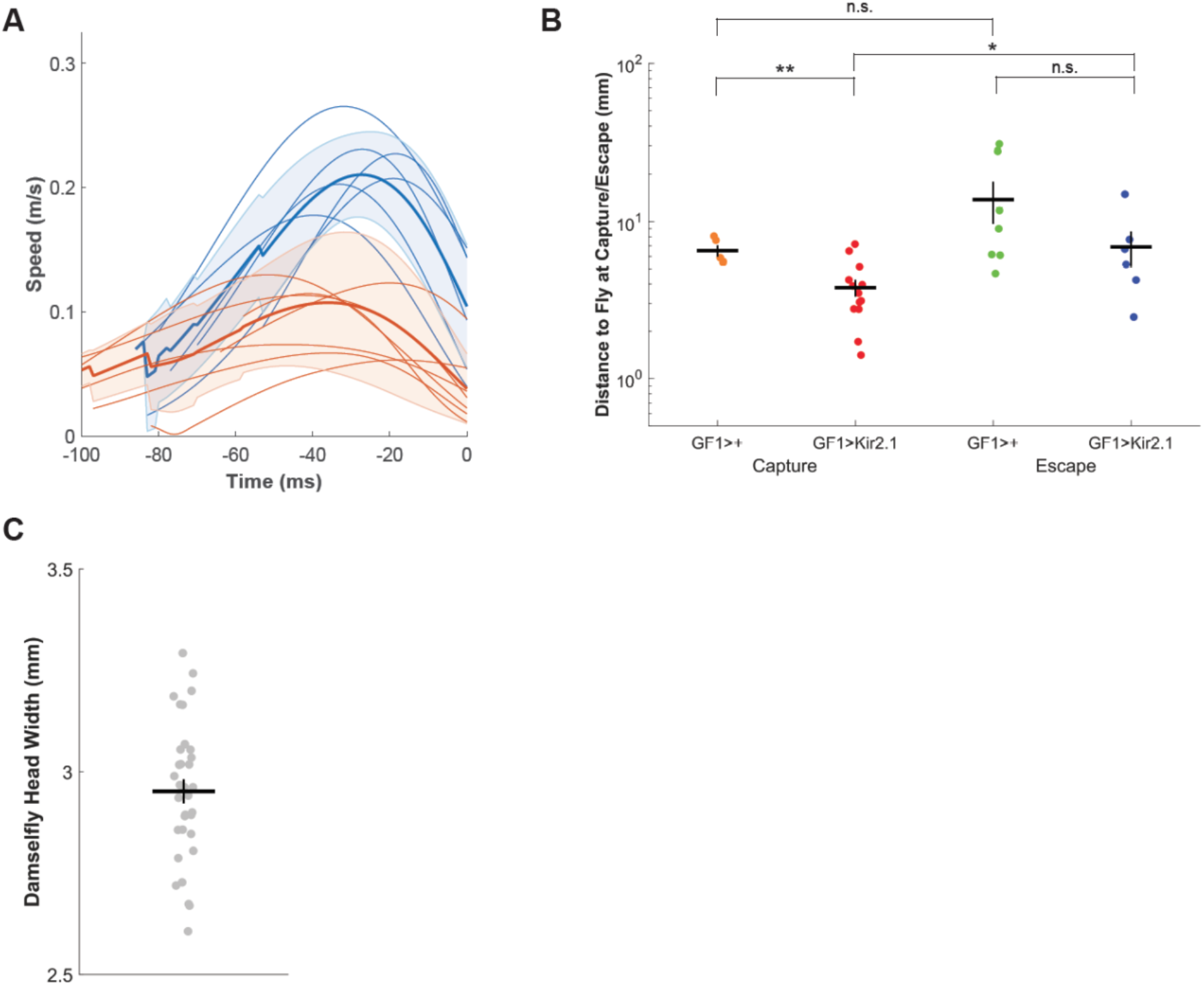
Damselfly attack kinematics additional analysis. Related to Figure 3. (A) Damselfly attack speed profiles for predation events ending in GF>Kir2.1 fly capture. Time = 0 represents the time of either contact or escape. Thin lines represent individual attack events that reach peak speeds above 0.13 m/s (blue) and below 0.13 m/s (red). Thick lines represent means and shaded regions represent standard deviations. (B) Distance between damselfly and fly at time of contact or escape. Colored dots represent individual predation events, black crosses represent mean ± SEM. GF1>+ Capture vs GF1>Kir2.1 Capture (**p=0.0047), GF1>+ Capture vs GF1>+ Escape (p=0.1759, n.s.), GF1>Kir2.1 Capture vs GF1>Kir2.1 Escape (*p=0.0356), GF1>+ Escape vs GF1>Kir2.1 Escape (p=0.1770, n.s.). p-values from two-sample t-test. (C) Distribution of damselfly head width measurements. n=33. Gray dots represent individual animals, black cross represents mean ± SEM

## REFERENCES

1. Hartline, D.K., and Colman, D.R. (2007). Rapid conduction and the evolution of giant axons and myelinated fibers. Curr Biol 17, R29–35. 10.1016/j.cub.2006.11.042.

2. Niven, J.E., Anderson, J.C., and Laughlin, S.B. (2007). Fly photoreceptors demonstrate energy-information trade-offs in neural coding. PLoS Biol 5, e116. 10.1371/journal.pbio.0050116.

3. Perge, J.A., Koch, K., Miller, R., Sterling, P., and Balasubramanian, V. (2009). How the optic nerve allocates space, energy capacity, and information. J Neurosci 29, 7917–7928. 10.1523/JNEUROSCI.5200-08.2009.

4. Sengupta, B., Faisal, A.A., Laughlin, S.B., and Niven, J.E. (2013). The effect of cell size and channel density on neuronal information encoding and energy efficiency. J Cereb Blood Flow Metab 33, 1465–1473. 10.1038/jcbfm.2013.103.

5. O ‘shea, M., Rowell, C.H.F., and Williams, J.L.D. (1974). The Anatomy of A Locust Visual Interneurone; the Descending Contralateral Movement Detector. Journal of Experimental Biology 60, 1–12. 10.1242/jeb.60.1.1.

6. Westin, J., Langberg, J.J., and Camhi, J.M. (1977). Responses of giant interneurons of the cockroachPeriplaneta americana to wind puffs of different directions and velocities. Journal of comparative physiology 121, 307–324. 10.1007/BF00613011.

7. Edwards, D.H., Heitler, W.J., and Krasne, F.B. (1999). Fifty years of a command neuron: the neurobiology of escape behavior in the crayfish. Trends Neurosci 22, 153–161. 10.1016/s0166-2236(98)01340-x.

8. Allen, M.J., Godenschwege, T.A., Tanouye, M.A., and Phelan, P. (2006). Making an escape: development and function of the Drosophila giant fibre system. Semin Cell Dev Biol 17, 31–41. 10.1016/j.semcdb.2005.11.011.

9. Fotowat, H., Harrison, R.R., and Gabbiani, F. (2011). Multiplexing of motor information in the discharge of a collision detecting neuron during escape behaviors. Neuron 69, 147–158. 10.1016/j.neuron.2010.12.007.

10. Kupfermann, I., and Weiss, K.R. (1978). The command neuron concept. Behavioral and Brain Sciences 1, 3–10. 10.1017/S0140525X00059057.

11. Eaton, R.C. (1983). Is the Mauthner Cell a Vertebrate Command Neuron? A Neuroethological Perspective on an Evolving Concept. In Advances in Vertebrate Neuroethology, J.-P. Ewert, R.R. Capranica, and D.J. Ingle, eds. (Springer US), pp. 629–636. 10.1007/978-1-4684-4412-4_31.

12. Schrameck, J.E. (1970). Crayfish swimming: alternating motor output and giant fiber activity. Science 169, 698–700. 10.1126/science.169.3946.698.

13. Eaton, R.C., Lavender, W.A., and Wieland, C.M. (1982). Alternative neural pathways initiate fast-start responses following lesions of the mauthner neuron in goldfish. Journal of comparative physiology 145, 485–496. 10.1007/BF00612814.

14. Kramer, A.P., and Krasne, F.B. (1984). Crayfish escape behavior: production of tailflips without giant fiber activity. Journal of Neurophysiology 52, 189–211. 10.1152/jn.1984.52.2.189.

15. Bacon, J.P., and Strausfeld, N.J. (1986). The dipteran ‘Giant fibre’ pathway: neurons and signals. Journal of Comparative Physiology A 158, 529–548. 10.1007/BF00603798.

16. Comer, C.M., Dowd, J.P., and Stubblefield, G.T. (1988). Escape responses following elimination of the giant interneuron pathway in the cockroach, Periplaneta americana. Brain Res 445, 370–375. 10.1016/0006-8993(88)91202-4.

17. Otis, T.S., and Gilly, W.F. (1990). Jet-propelled escape in the squid Loligo opalescens: concerted control by giant and non-giant motor axon pathways. Proc Natl Acad Sci U S A 87, 2911–2915. 10.1073/pnas.87.8.2911.

18. von Reyn, C.R., Breads, P., Peek, M.Y., Zheng, G.Z., Williamson, W.R., Yee, A.L., Leonardo, A., and Card, G.M. (2014). A spike-timing mechanism for action selection. Nat Neurosci 17, 962–970. 10.1038/nn.3741.

19. Richardson, J.M.L., and Baker, R.L. (1997). Effect of Body Size and Feeding on Fecundity in the Damselfly Ischnura verticalis (Odonata: Coenagrionidae). Oikos 79, 477. 10.2307/3546890.

20. Woodward, G. (2001). Dragonflies: Behaviour and Ecology of Odonata. Freshwater Biology 46, 141–143. 10.1111/j.1365-2427.2001.00664.x.

21. Baines, R.A., Uhler, J.P., Thompson, A., Sweeney, S.T., and Bate, M. (2001). Altered electrical properties in Drosophila neurons developing without synaptic transmission. J Neurosci 21, 1523–1531. 10.1523/JNEUROSCI.21-05-01523.2001.

22. Luan, H., Peabody, N.C., Vinson, C.R., and White, B.H. (2006). Refined spatial manipulation of neuronal function by combinatorial restriction of transgene expression. Neuron 52, 425–436. 10.1016/j.neuron.2006.08.028.

23. Pfeiffer, B.D., Ngo, T.T., Hibbard, K.L., Murphy, C., Jenett, A., Truman, J.W., and Rubin, G.M. (2010). Refinement of tools for targeted gene expression in Drosophila. Genetics 186, 735–755. 10.1534/genetics.110.119917.

24. Williamson, W.R., Peek, M.Y., Breads, P., Coop, B., and Card, G.M. (2018). Tools for Rapid High-Resolution Behavioral Phenotyping of Automatically Isolated Drosophila. Cell Rep 25, 1636–1649 e1635. 10.1016/j.celrep.2018.10.048.

25. Card, G., and Dickinson, M. (2008). Performance trade-offs in the flight initiation of Drosophila. J Exp Biol 211, 341–353. 10.1242/jeb.012682.

26. Klapoetke, N.C., Nern, A., Peek, M.Y., Rogers, E.M., Breads, P., Rubin, G.M., Reiser, M.B., and Card, G.M. (2017). Ultra-selective looming detection from radial motion opponency. Nature 551, 237–241. 10.1038/nature24626.

27. von Reyn, C.R., Nern, A., Williamson, W.R., Breads, P., Wu, M., Namiki, S., and Card, G.M. (2017). Feature Integration Drives Probabilistic Behavior in the Drosophila Escape Response. Neuron 94, 1190–1204 e1196. 10.1016/j.neuron.2017.05.036.

28. Ache, J.M., Polsky, J., Alghailani, S., Parekh, R., Breads, P., Peek, M.Y., Bock, D.D., von Reyn, C.R., and Card, G.M. (2019). Neural Basis for Looming Size and Velocity Encoding in the Drosophila Giant Fiber Escape Pathway. Curr Biol 29, 1073–1081 e1074. 10.1016/j.cub.2019.01.079.

29. Eaton, R.C., Nissanov, J., and Wieland, C.M. (1984). Differential activation of Mauthner and non-Mauthner startle circuits in the zebrafish: Implications for functional substitution. Journal of Comparative Physiology A 155, 813–820. 10.1007/BF00611598.

30. Herberholz, J., Sen, M.M., and Edwards, D.H. (2004). Escape behavior and escape circuit activation in juvenile crayfish during prey-predator interactions. J Exp Biol 207, 1855–1863. 10.1242/jeb.00992.

31. Hecker, A., Schulze, W., Oster, J., Richter, D.O., and Schuster, S. (2020). Removing a single neuron in a vertebrate brain forever abolishes an essential behavior. Proceedings of the National Academy of Sciences 117, 3254-3260. doi:10.1073/pnas.1918578117.

32. Hodgkin, A.L., and Huxley, A.F. (1952). A quantitative description of membrane current and its application to conduction and excitation in nerve. J Physiol 117, 500–544. 10.1113/jphysiol.1952.sp004764.

33. Luan, H., Diao, F., Scott, R.L., and White, B.H. (2020). The Drosophila Split Gal4 System for Neural Circuit Mapping. Front Neural Circuits 14, 603397. 10.3389/fncir.2020.603397.

34. Gabbiani, F., Krapp, H.G., and Laurent, G. (1999). Computation of object approach by a wide-field, motion-sensitive neuron. J Neurosci 19, 1122–1141. 10.1523/jneurosci.19-03-01122.1999.

35. Hedrick, T.L. (2008). Software techniques for two- and three-dimensional kinematic measurements of biological and biomimetic systems. Bioinspir Biomim 3, 034001. 10.1088/1748-3182/3/3/034001.

